# PRL-1 is required for neuroprotection against olfactory CO_2_ stimulation in Drosophila

**DOI:** 10.1101/355438

**Authors:** Pengfei Guo, Xiao Xu, Fang Wang, Xin Yuan, Yinqi Tu, Bei Zhang, Huimei Zheng, Danqing Yu, Wanzhong Ge, Zhefeng Gong, Weiqiang Gao, Xiaohang Yang, Yongmei Xi

## Abstract

The Mammalian phosphatase of regenerating liver (PRL) family is primarily recognized for its oncogenic properties. Here we found that in Drosophila, loss of *prl-1* resulted in CO2-induced brain disorder presented as irreversible wing hold up with enhancement of Ca2+ responses at the neuron synaptic terminals. Overexpression of Prl-1 in the nervous system could rescue the mutant phenotype. We show that Prl-1 is particularly expressed in CO2-responsive neural circuit and the higher brain centers. Ablation of the CO2 olfactory receptor, Gr21a, suppressed the mutant phenotype, suggesting that CO2 acts as a neuropathological substrate in absence of Prl-1. Further studies found that the wing hold up is an obvious consequence upon knockdown of Uex, a magnesium transporter, which directly interacts with Prl-1. Conditional expression of Uex in the nervous system could rescue the phenotype of *prl-1* mutants. We demonstrate that Uex acts genetically downstream of Prl-1. Our findings provide important insights into mechanisms of Prl-1 protection against olfactory CO2 stimulation induced brain disorder at the level of detailed neural circuits and functional molecular connections.

## Background

Drosophila phosphatase of regenerating liver-1(prl-1) is the only homologous gene of the mammalian PRL family (including PRL-1, PRL-2 and PRL-3) that belongs to the smallest class (molecular masses of 20-22kDa) of protein tyrosine phosphatases (PTPs) (Diamond, Cressman et al., 1994, Zeng, Hong et al., 1998). In recent years, much attention has been paid to PRLs due to their implication in various cancers (Bessette, Qiu et al., 2008, Rios, Li et al., 2013, Saha, Bardelli et al., 2001, Zeng, Dong et al., 2003). The highly expressing PRL-3 has almost been exclusively and particularly correlated to disease aggressiveness and clinical outcome for such multiple tumor types as colorectal(Molleví, Aytes et al., 2008, Xing, Peng et al., 2009), ovarian(Polato, Codegoni et al., 2005, Ren, Jiang et al., 2009), breast(Wang, Peng et al., 2006) and gastric(Wang, Cai et al., 2009) cancers. Analysis of PRL-1, PRL-2 mRNA in a large number of variant human tissue specimens revealed its significant overexpression in hepatocellular and gastric carcinomas, but significant underexpression in ovarian, breast, kidney carcinomas (Dumaual, Sandusky et al., 2012). Variant expression was also noted in other non-cancer tissue types(Dumaual, Sandusky et al., 2006). These contradictory results suggested high tissue-specific and a pleiotropic role for PRL in the diseases process. Nevertheless, neither the mechanism nor the regulation of PRL have been clarified, the exact biological function of these PRL molecules remains currently unknown and a clear mutant background of PRL has yet to be established in any genetic model.

Characterization of the Prl family members has been conducted for early embryos of *Drosophila*, amphioxus and zebrafish, where Prls were seen to be uniformly expressed in the central nervous system (CNS) and suggested to be likely involved in early neural development (Lin, Lee et al., 2013). In a mouse model PRL-2 was noted as ubiquitously expressed, particularly in the hippocampal pyramidal neurons, ependymal cells, cone and rod photoreceptor cells (Gungabeesoon, Tremblay et al., 2016). We initially investigated the function of *Drosophila* Prl-1 by generation of *prl-1* mutant using CRISPR/Cas9 method (Bassett & Liu, 2014, Bassett, Tibbit et al., 2013). Surprisingly, we found that in the absence of Prl-1, a CO_2_-induced irreversible wing hold up phenotype resembling spasticity develops. This led us to discover a possible role for Prl-1 in the nervous system.

CO_2_-evoked behavioral responses in winged insects are important for food foraging, reproduction and survival (Guerenstein, Christensen et al., 2004, McMeniman, Corfas et al., 2014, Stange & Stowe, 2015). *Drosophila*, in particular, is highly sensitive to CO_2_ where the detection of a CO_2_ threshold is usually accompanied by clear physiological and behavioral responses. These have been previously explained in terms of anesthetic and toxic effects related to high concentrations of CO_2_ (Badre, Martin et al., 2005, Dijken, Sambeek et al., 1977), or as a stress odorant eliciting avoidance behavior upon the detection of CO_2_ levels as low as 0.1% (Suh, Wong et al., 2004). The recent discovery that *Drosophila* is equipped with a single population of CO_2_-responsive neurons harbored in the third segment of the antenna and that the avoidance behavior is mediated by two chemosensory receptors, Gr21a and Gr63a, represents a significant breakthrough in understanding how *Drosophila* senses and processes CO_2_ stimulus (Jones, Cayirlioglu et al., 2007, Scott, Jr et al., 2001, Suh et al., 2004). Upon CO_2_ stimulation, avoidance behavior can be assessed using a T-maze assay in laboratory conditions. Correspondingly, the neural activity at sensory neuron synaptic terminals in the CO_2_-responsive V- glomeruli of the antennal Lob (AL) can be detected by Ca^2+^ imaging (Jones et al., 2007, Suh et al., 2004). However, to what extent CO_2_ stimuli has influence from the level of functional molecular connections at that of detailed neural circuits to the physiological function at bodily levels remains obscure. Here we reveal the localization of Prl-1 in CO_2_-responsive neural circuit and the higher brain centers and uncover a Prl-1 involved protective mechanism, based on its interaction with the Uex in the nervous system.

Uex, is the homologous gene of human ancient domain proteins (ACDPs), also known as CNNMs including CNNM1-4, of which CNNM4 acts as a Na^+^/Mg^2+^ exchanger and regulates Mg^2+^ efflux (Yamazaki, Funato et al., 2013). CNNMs were identified to interact with hPRL-1 or hPRL-2 in the mediation of cancer metastasis (Funato, Yamazaki et al., 2014, Hardy, Uetani et al., 2015).A recent study on human genetic diseases has reported that mutations in CNNM2 are causative for seizures and mental retardation in patients with hypomagnesemia(Arjona, de Baaij et al., 2014). In this study we initially confirmed that their homologues, Prl-1 and Uex, also physically interacted in Drosophila. We show that the wing hold up is an obvious consequence upon knockdown of Uex and the expression of Uex is dramatically decreased in *prl-1* mutants. From the previous focus being almost exclusively oncogenic, it was surprising to uncover a highly significant Prl-1-Uex complex based neuroprotective role in *Drosophila*. Ectopic expression of either Drosophila Prl-1 or hPRL in the nervous system could rescue the mutant phenotype. Our study may open up a potential new focus upon the functional pathway of the hPRL beyond studies of oncogenic properties.

## Results

### Loss of Prl-1 results in irreversible wing hold up in *Drosophila*

An often overlooked behavioral phenomenon when using standard CO_2_ anesthesia in *Drosophila* is that the flies respond with a temporary holding up of wings which ceases upon recovery. Surprisingly, we observed an irreversible wing hold up phenotype developing in *prl-1* mutants beyond CO_2_ stimulation. To characterize the occurrence of this wing hold up phenotype, we began by dividing the *prl-1* mutant or wild type flies into several age groups (day 1, day 2, or day 3), keeping them separately (n=180 for each group, 20 flies per vial). All experimental groups were subjected to acute CO_2_ under a flow of 5L/m for 20 sec as a minimum manipulation. Non CO_2_-treated groups served as controls. After recovery, all age groups of WT flies showed normal wing posture. However, about 85% of mutants that had received the CO_2_ administration on day 3 showed irreversible wing hold up (Fig.1A-D and Fig. S1B). The mutants that had either received, or had not received the specific CO_2_ administration on day 1, began to show latent and progressive wing hold up, gradually reaching a percentage of 60% around day-21. This was both a considerably lower level and a slower response than that of the flies that had received the CO_2_ stimulation on day 2 or day 3 (Fig. S1B). These observations revealed that the 3-day old mutant flies were particularly sensitive to CO_2_ stimulation and present a rapid prevalence of irreversible wing hold up. As male *prl-1* mutants displayed the more prominent phenotype upon the CO_2_ stimulation (Fig. S1A), we exclusively analyzed male responses in the following experiments.

**Fig. 1.**
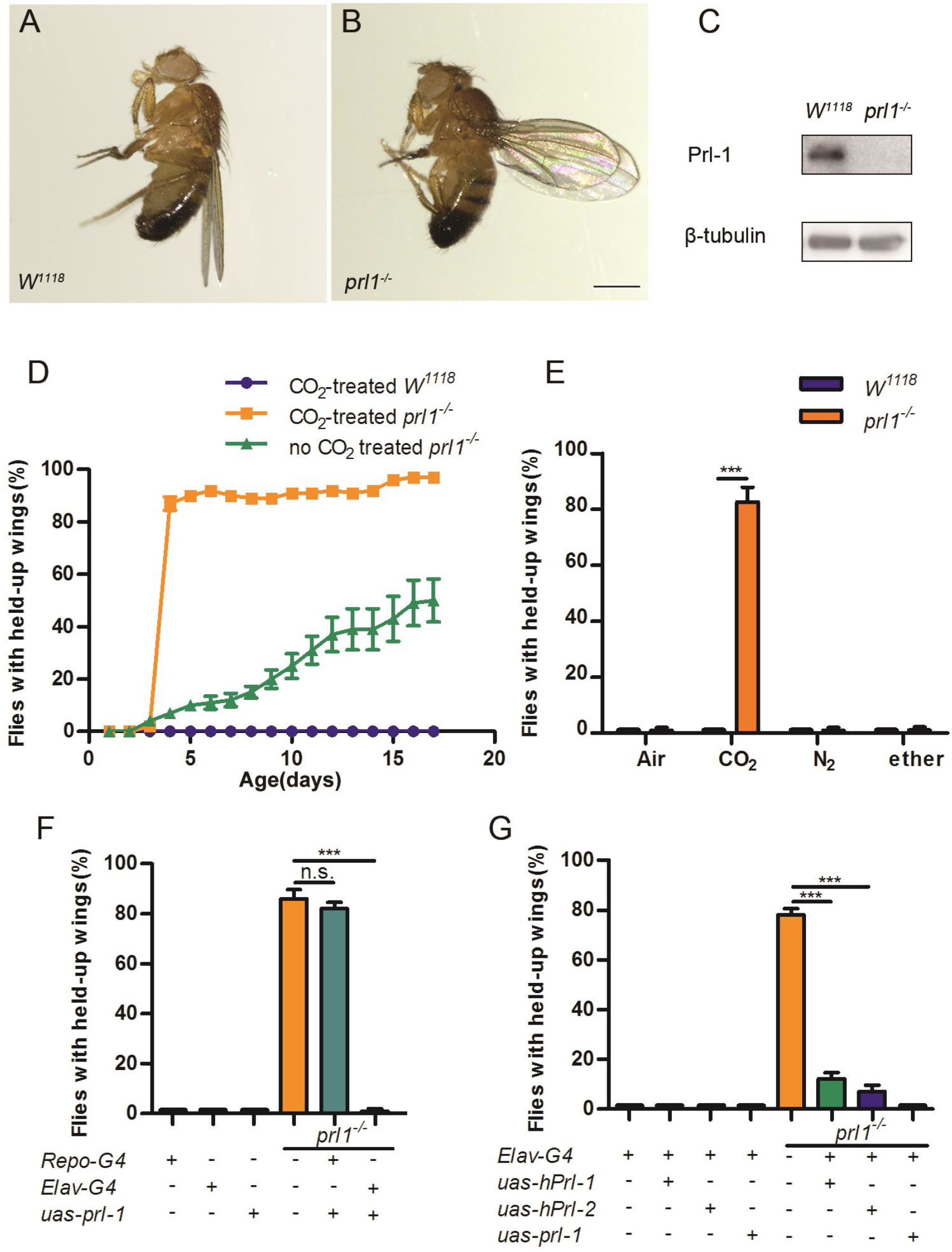
Loss of Prl-1 results in CO_2_-induced wing hold up in *Drosophila*. **(A-B)** Wild type controls and *prl-1* mutant animals with exposure of high concentrations of CO_2_. *prl1*^−/−^ denotes the *prl-1* mutant allele. *prl-1* mutant animals display wing hold-up. **(C)** Western blot on adult fly extracted from wild type and *prl-1* mutant animals. Lysates were probed with anti-Prl-1 and anti-Tubulin antibodies. **(D)** *prl-1* mutant animals display a quick response of wing hold-up phenotype with exposure to CO_2_, while wild type flies still hold wings normally. The mutants also show an age-dependent increase of held-up wings when kept in the normal CO_2_ levels of the ambient environment. **(E)** The response of *prl-1* mutant flies to alternative anesthetics, carbon dioxide (CO_2_), ether, and nitrogen (N_2_). **(F)** Prl-1 expression from a *Uas-prl-1* transgene with two different drivers (*Repo-Gal4* or *Elav-Gal4*) was used to rescue the abnormal wing posture in *prl-1* mutants. Only *Elav-gal4* could entirely recover the normal wing posture. **(G)** Expression of human *Prl*s (*hPrl-1* or *hPrl-2*) in *prl-1* mutant background driven by *Elav-Gal4* could rescue the wing hold-up phenotype. Statistics for **(D-G)**: results are plotted as means ± SEM. Two-tailed Student’s t test with ***p<0.001, n=100 for all groups. Scale bar: 200x pix in B.

We then considered whether this wing hold up in *prl-1* mutants had resulted from a side-effect of general anesthesia. Gaseous nitrogen (N_2_) or volatile ether (Dijken et al., 1977, Van Voorhies, 2009) was used to anesthetize 3 day old *prl-1* mutant flies, with the CO_2_ treatment used as the control. After recovery, neither N_2_ nor ether anesthesia groups displayed the wing hold up as seen in the CO_2_ treated groups (Fig. 1E). These observations indicated that irreversible wing hold up, as induced by CO_2_ exposure in *prl-1* mutants, does not simply occur as a direct result of general anesthesia.

### Expressing Prl-1 or hPRL in the nervous system can rescue the mutant phenotype

We next conducted a series of assays to screen the specific tissues or cells of *prl-1* mutants with overexpression of Prl-1, in which the wing hold up could be rescued. The *uas-prl-1* transgene line was expressed in the mutant flies under the control of several tissue or cell type-specific Gal4 drivers, including Tublin-gal4, pan-neuronal Elav-Gal4, Repo-Gal4, TH-Gal4, motor neuronal D42-Gal4, and muscle Mhc-Gal4. Surprisingly, among all drivers genetically manipulated in the prl-1 mutants, only the pan-neuronal expressed Elav-Gal4 could completely rescue the wing hold up in the 3-day old groups that had either received, or not received the acute CO_2_ treatment (Fig.1F and Fig. S1C). While other drivers, such as D42-Gal4 exhibited a partial rescue, the Tublin-gal4, TH-Gal4, Mhc-gal4 drivers failed to provide any rescue of the mutant phenotype (Table S1). These observations indicated a potent role for Prl-1 in the *Drosophila* nervous system, specifically relating to CO_2_ stimulation. Given this neurologic impairment in *prl-1* mutant flies, we then tested the function of hPRL in the context of the *Drosophila* brain. We generated transgenic flies containing hPRL-1 or hPRL-2. Ectopic expression of either UAS-hPRL-1 or UAS-hPRL-2 driven by Elav-Gal4 was able to rescue the wing hold up (Fig. 1G). Our results suggest that hPRL may have retained this conserved neuroprotective function.

### Prl-1 is enriched in the CO_2_ neural circuit and higher brain centers

To evaluate the expression pattern of Prl-1 in the brain, we first performed an immunoblot assay with the whole head tissues of adult flies. A positive signal was detected in the control animals by using a rabbit polyclonal antibody against the full-length peptide of Prl-1, which was abolished in *prl-1* mutants (Fig. S1D). To accurately target the Prl-1 expressing neurons, we also generated a transgenic Gal4 line which contained a 6.1Kbp of genomic DNA immediately upstream to the open reading frame of the *prl-1* gene. We flanked the Gal4 sequence with both the 5’ and 3’ flanking regions of the *prl-1* gene and constructed a fused green fluorescent protein (EGFP) with Prl-1 sequence in the N-terminal to make a high-fidelity reporter. By directly viewing GFP fluorescence in the transgenic flies expressing EGFP-Prl-1 under the control of the Prl-Gal4 driver, the whole head showed green with particularly robust GFP signals in the third segment of antennae (Fig.2A-A’). Confocal scanning revealed that the Prl-1 protein was apparently expressed in the basiconic sensillum (Fig.2B-B’), which houses CO_2_-responsive neurons (Scott et al., 2001, Suh et al., 2004). Prl-1 expression had also expanded along the axons of the olfactory CO_2_ neurons to its projected V-glomeruli in the antennal lobe (AL), and further to the higher processing center, the mushroom body (MB) (Kwon, Dahanukar et al., 2007, Suh et al., 2004). The distribution of EGFP-Prl-1 in the AL and the MB was also assessed using immunofluorescence staining of the adult brain with a GFP antibody (Fig.2C-D’). It is notable that the enrichment of Prl-1-Gal4 expression in the stereotyped V-glomeruli corresponds very well to the identical dendritic structure of CO_2_ sensory neurons expressing Gr21a-Gal4 and Gr63a-Gal4 (Jones et al., 2007) and their projection neurons expressing PNv-Gal4s (v201089, v200516) (Lin, Chu et al., 2013) (Fig.3A, and Fig. S2).

**Fig. 2.**
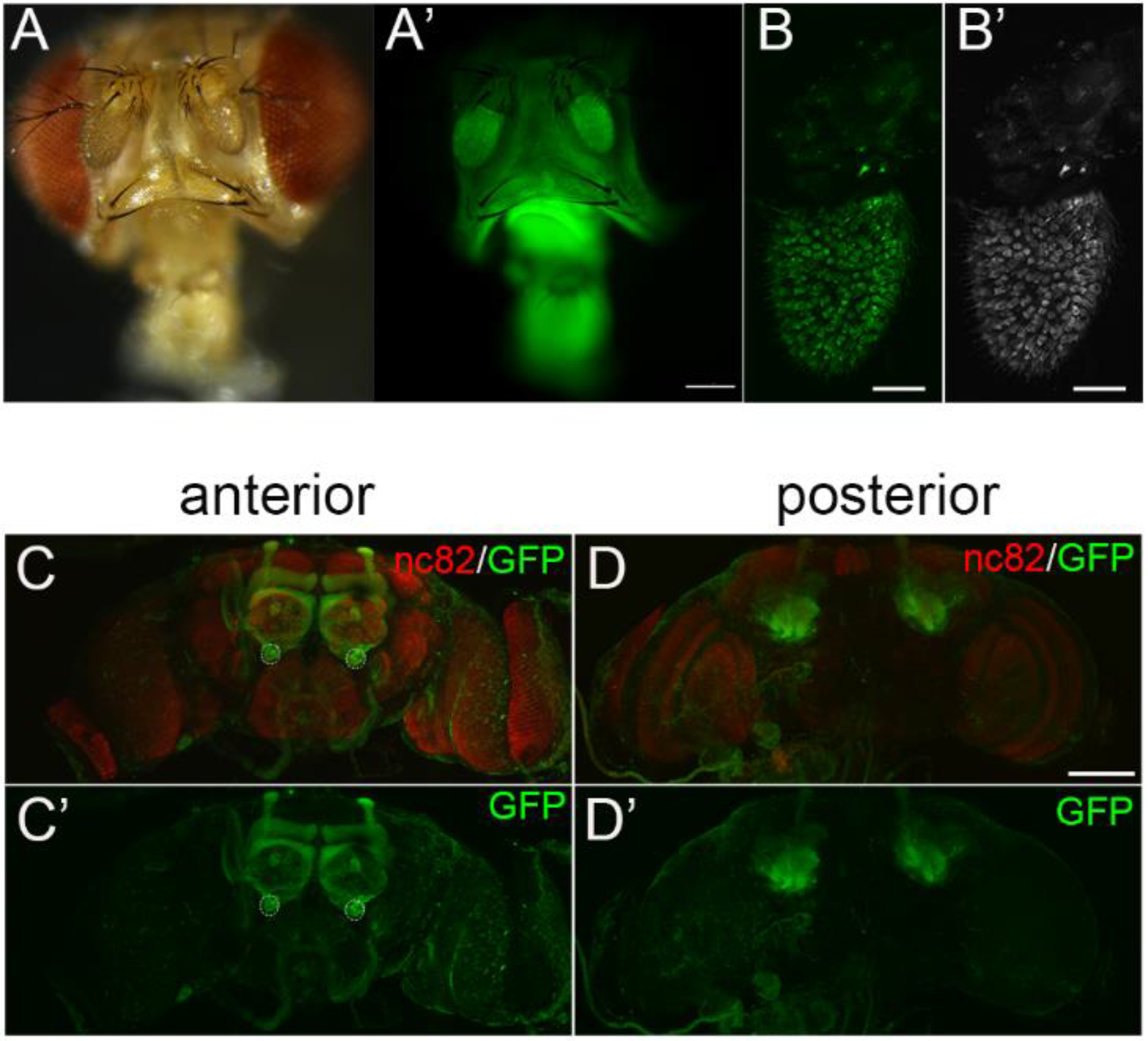
Prl-1 is enriched in the CO_2_ neural circuit and higher brain centers. **(A-A’)** Direct GFP fluorescence of *EGFP-prl-1* with *prl-Gal4* driver. **(B-B’)** Confocal scanning of the antennae shows the distribution of Prl-1 in the region. **(C-D’)** Immunofluorescence staining reveals the spatial expression of UAS-EGFP-Prl-1 with Prl-Gal4 driver line in the adult brain. Dotted circles label the V-glomeruli. The adult brains were stained with anti-Brp^nc82^(red)and anti-GFP (green). Scale bar:100μm in **(D)**, 200x pix in A’ and B’.

### Ablation of Gr21a, an olfactory CO_2_ receptor, suppresses the *prl-1* mutant phenotype

Analysis of CO_2_-evoked avoidance responses using T-maze assays (Kwon et al., 2007, Suh et al., 2004) revealed no significant differences between *prl-1* mutants and the WT (Fig. S3A). This indicates that the olfactory sensing of CO_2_ remains active in *prl-1* mutants. However, irreversible wing hold up in the *prl-1* mutants proceeded with age when maintained in the ambient environment, and at a highly accelerated rates for 3-day old flies treated with the acute CO_2_ stimulation. We then considered whether the irreversible wing hold up could be blocked by suppressing CO_2_ perception via removing ligands from the receptors. We employed the RNA interference system to genetically knock down Gr21a in the CO_2_ sensory neurons. The bigeneric progeny of *prl-1* mutants, each group bearing Elav-Gal4 or Gr63a-Gal4 along with Gr21a-RNAi, no longer displayed any rapid prevalence of irreversible wing hold up under acute CO_2_ exposure for 3-day old flies. Progressive irreversible wing hold up had also dramatically decreased from 60% to less than 5% in the non CO_2_ treated groups (Fig.3B and Fig.S3B). However, driven by Gr63a-Gal4, overexpression of Prl-1 only in the CO_2_ sensory neurons could not rescue the phenotype (data not shown). These observations suggest that the mutant flies are highly susceptible to brain disorder when the Prl-1 linked defense against CO_2_ insult is compromised. Therefore, the irreversible wing hold up is a deficit in olfactory information processing in the central brain, where CO_2_ might act as a neuropathological trigger in absence of Prl-1.

**Fig. 3.**
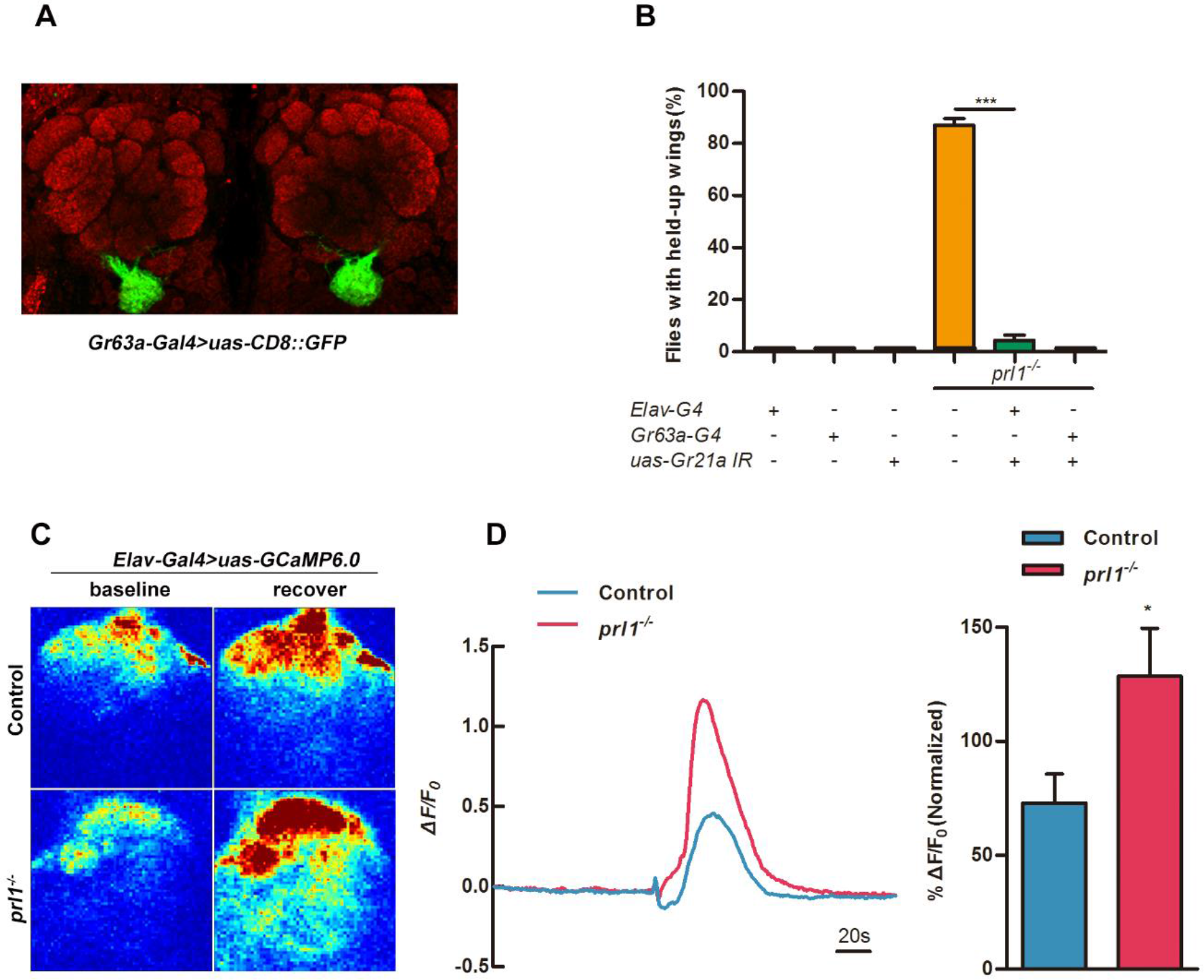
Ablation of Gr21a, an olfactory CO_2_ receptor, suppresses the phenotype of the mutants. **(A)** Olfactory neurons expressing Gr63a-Gal4; UAS-CD8::GFP (green) and anti-Brp^nc82^ (red) of the V glomeruli in the antennal lobe. **(B)** In *prl-1* mutant background, genetic ablation of CO_2_ sensory neurons driven by *Elav-Gal4* or *Gr63a-Gal4* could inhibit the flies to avoid wing hold-up. **(C-D)** Calcium responses in the antennal lobe with CO_2_ stimulation for control or *prl-1* mutant flies. **(A)** Representative images of GCaMP6.0 fluorescence (%ΔF/F) obtained from in vivo functional imaging at baseline and after CO_2_ stimuli to control or *prl-1*mutant flies. **(B)** GCaMP6.0 responses from control or *prl-1* mutant flies across CO_2_ presentation with bar graph quantitation. Statistics for **(D)** and **(F)**: results are plotted as means ± SEM. Two-tailed Student’s t test with *p<0.05, n=7 for each group.

### Enhancement of CO_2_-evoked Ca^2+^ activity and elevated ROS in the *prl-1* mutants

Upon activation of CO_2_-responsive neural circuit driven aversion responses, Ca^2+^ transients could be visualized using two-photon microscopy with the expression of calcium-sensitive fluorescent protein (GCaMP) in the AL (Jones et al., 2007, Suh et al., 2004). We analyzed the V-glomerular activation pattern by Ca^2+^ imaging in WT and *prl-1* mutants upon CO_2_ -evoked aversion responses. The GCaMP indicator (UAS-GCaMP6.0) is driven by the Elav-GAL4 activator in all neurons (Jones et al., 2007, Wang, Wong et al., 2003). Upon 20 sec CO_2_ exposure, the V glomeruli were activated (Fig.3C). The intensity of Ca^2+^ activity viewed in the *prl-1* mutants was 2-fold higher than that in the controls (average peak ΔF/F of *prl*-1^−/−^ is 1.21±0.51; average peak ΔF/F of WT is 0.55±0.46) (Fig. 3D-E). The overexpression of a *uas-prl-1* transgene driven by Elav-Gal4 in the mutants could restore CO_2_-evoked Ca^2+^ activity to their original levels (Fig.S3D-F).

Oxidative stress represents a number of sequential and integrated processes that lead to cell vulnerability in the brain (Coyle & Puttfarcken, 1993).We took advantage of the GSTD1-ARE-GFP flies to evaluate the reactive oxygen species (ROS) levels in the *prl-1* mutants, where GSTD1 is regulated via the Keap1/cnc signaling pathway in response to oxidative stress (Nguyen, Nioi et al., 2009, Sykiotis & Bohmann, 2008). By viewing GFP fluorescence in different age groups of flies, higher levels of ROS, as indicated by stronger GFP signals, were observed in *prl-1* mutants as compared to those of the control animals (Fig. S3C). We also performed knockdown of the cnc levels in PRL-1 mutants, resulting in exacerbated wing hold up phenotype in the mutants (data not shown).The pan-neuronal Elav-gal4 driven Prl-1 overexpression was able to rescue the elevated ROS phenotype (Fig. S3C). This data may serve as an indicator that PRL-1 may function as a neurological antioxidant via the Keap1/cnc pathway.

### Irreversible wing hold up results from a decrease of Uex in the *prl-1* mutants

We next asked whether *prl-1* cases the underlying genes responsible for the irreversible wing hold up. We focused on *Drosophila* Uex, the only known homologous gene of the CNNM family members that have been recognized as Prl-binding partners in mammalian cells (Funato et al., 2014, Hardy et al., 2015). To examine whether Prl-1 also interacts with Uex in *Drosophila*, WT Prl-1 and catalytically inactive mutant Prl-1-D77A/C109S constructs were generated with HA-tagged at their N-terminus. These were transiently expressed in S2 cells and the lysates were subjected to IP with an anti-HA antibody. Co-immunoprecipitation of endogenous Uex could be detected when exogenous HA-Prl-1 was precipitated from S2 cells (Fig. 4A) revealing that Uex can also interact with Prl-1 in *Drosophila*. A GST-pulldown assay validated the direct interaction between Prl-1 and Uex, whereas this interaction was decreased with the mutation of the catalytic sites Prl-1-C109S or Prl-1-D77A/C109S (Fig. 4B). The Co-IP results showed that these amino acid substitution mutants of Prl-1 largely decreased the Prl-1-Uex binding ability (Fig. 4A). This suggested that Uex does not function as a phosphate substrate of Prl-1. The co-localization of Prl-1 and Uex on the plasma membrane of S2 cells was also detected by double-immunofluorescence staining with the antibodies against Prl-1 and Uex (Fig. 4C).

**Fig. 4.**
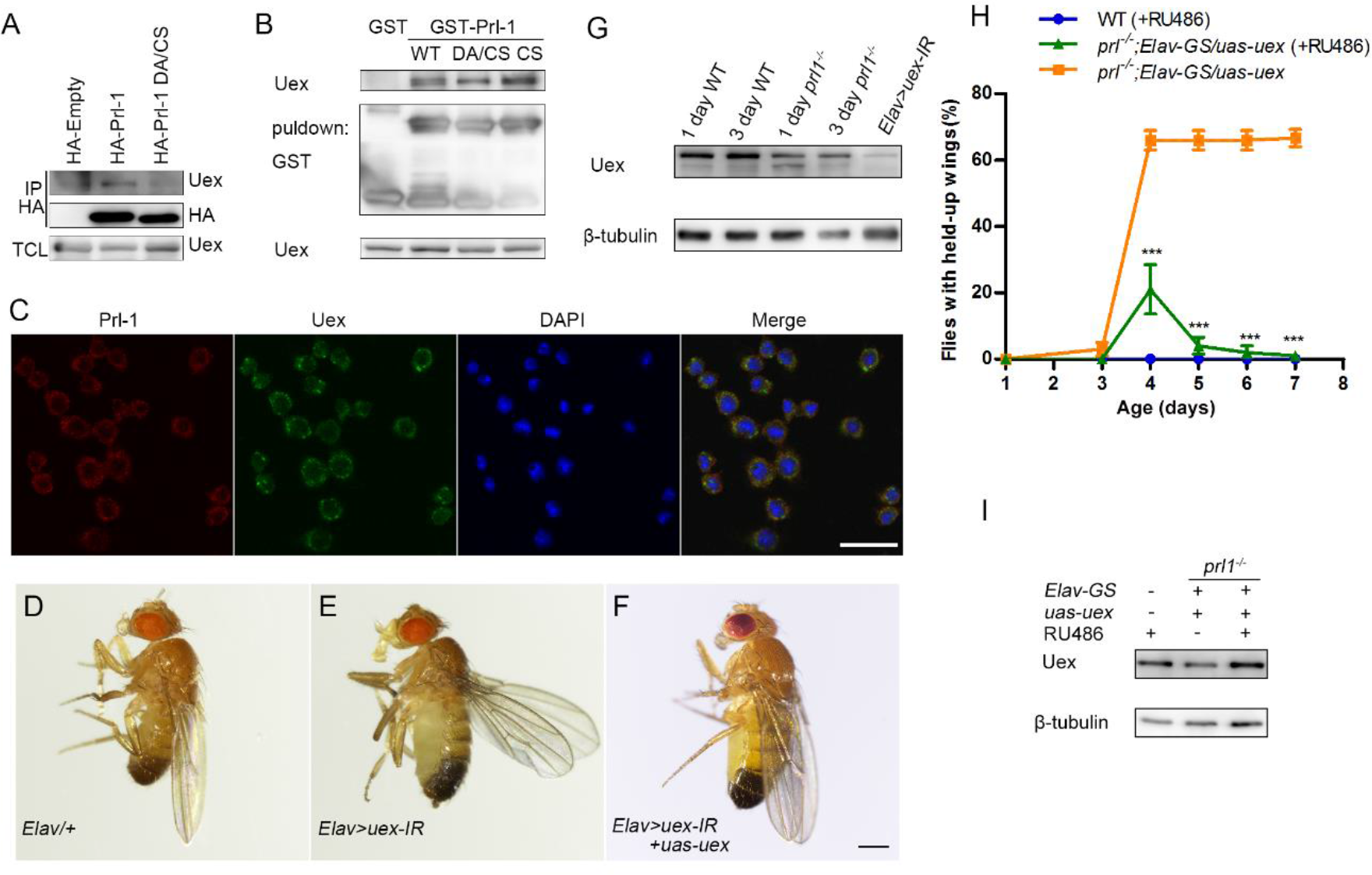
Wing hold up phenotype results from a decrease of Uex in the *prl-1* mutants. **(A)** S2 cells were transiently transfected with either empty or HA-tagged Prl-1(WT or D77A/C109Smutant). Lysates were subjected to immunoprecipitates and immunoblot analyses with the anti-HA and anti-Uex antibodies. **(B)** Lysates were extracted from WT adult flies for GST pulldown assay with purified GST-tagged Prl-1(WT, C109S or D77A/C109S) proteins. **(C)** S2 cells were subjected to immunofluorescence staining with anti-Prl-1(red), anti-Uex(green), and DAPI (blue). **(D-F)** Loss of Uex function in fly nervous system results in abnormal wing posture.**(D)** Adult male flies expressing the following transgenic lines under the control of Elav-Gal4.**(E)** *uas-uex* RNAi (*uex-IR*), knockdown of Uex driven by Elav-Gal4 caused the wing hold-up phenotype (Genotype: *Elav/*+*; uas-uex-IR/*+).**(F)** Held-up wings induced by Uex can be restored to normal posture with *uas-uex* expression (Genotype: *Elav-Gal4/*+*; uas-uex-IR/uas-uex*), as in wild type controls (Genotype: *Elav-Gal4/*+). **(G)** Western blot analyzes protein level of Uex in 1-day and 3-day oldcontrol and *prl-1* mutant flies.**(H)** Conditionally increasing expression of Uex in *prl-1* mutant flies. Expression of Uex was induced with 0.5mg/ml RU486 in experimental groups for 2 days, and all flies were exposed with 20s CO_2_ on day 3.**(I)** Western blot analysis of Uex protein of 5-day old control, *prl-1* mutant, and GeneSwitch-induced *prl-1* mutant flies by feeding with 0.5mg/ml RU486. Scale bar.

Given the abnormal wing posture in *prl-1* mutant flies and a direct interaction existing between Prl-1 and Uex, we then searched for a similar phenotype of abnormal expression of Uex. However, the loss of Uex function resulting from CRISPR/Cas9 led to embryonic lethality. RNA interference was then used to knock down Uex expression in adult flies and a panel of Gal4 lines was employed to detect wing posture. Strikingly, all flies expressing Uex-RNAi (line 36116) driven by pan-neuronal Elav-Gal4 displayed a similar irreversible wing hold up, 1 day after eclosion (Fig. 4E and Fig.S4). This was associated with short life span of no more than 10 days. Several Gal4 lines were employed to map abnormal wing posture resulting from the knockdown of Uex. Among all these differing cell-type-or region-specific Gal4 lines, expressed within CNS or peripheral nervous system (PNS) or muscles, only Elav-Gal4 could induce the irreversible wing hold up (Fig. S4). By analyzing the pan-neuronal RNAi-induced knockdown of Uex expression, our Western blot data showed that a single protein with a mass of ~92 kDa had been obviously reduced compared to the wild type (Fig. 4G). We also generated a UAS-Uex line using the complete protein coding sequence to exclude any off-target effect. As figure 4F shows, these flies displayed normal wings with co-expression of UAS-Uex and Uex-RNAi lines. These results indicate that irreversible wing hold up is directly correlated to the function of Uex, the knockdown of Uex in the nervous system resulting in the same phenotype.

We hypothesize that the irreversible wing hold up might be accounted for by the decreased expression of Uex in the *prl-1* mutants, considering the direct interaction between Prl-1 and Uex in the brain. We probed the rate of Uex expression within the different age groups of flies after eclosion. Compared with the control animals, the protein level of Uex had decreased in *prl-1* mutant flies (Fig. 4G). More strikingly, there was a gradual degradation over time of Uex protein, from the newly eclosed mutant flies to the 3 day old mutant flies (Fig. 4G). This may explain why a particularly rapid prevalence of wing hold up occurs for 3-day old mutant flies treated with acute CO_2_. Alternatively, we performed ectopic expression of Uex in the *prl-1* mutant background to test whether Uex functions downstream of Prl-1 signaling. A conditional RU486-dependent Gal4 (GeneSwitch) was used to induce tissue-specific transgene expression (Osterwalder, Yoon et al., 2001). As RU486 was fed to the flies at the adult stage, *prl-1* mutant flies that were exposed with 20s CO_2_ on day 3, showed a reversible wing posture within one week (Fig. 4H). The RU486-induced Uex overexpression in *prl-1* mutants was verified by Western blotting (Fig. 4I). The Uex protein was recovered to a high level using the Elav-GeneSwitch driver. These results confirmed that Prl-1 positively regulates Uex at the genetic and molecular level in *Drosophila*.

## Discussion

Here, we have identified in a *Drosophila* model that, in the absence of Prl-1, olfactory CO_2_ stimulation can cause irreversible wing hold up, negating any possibility of flight. The *prl-1* mutant flies retain normal responses to anesthesia including CO_2_, N_2_ or volatile ether and also remain responsive to CO_2_ with no significant change in avoidance behavior. It seems that the sensing of CO_2_ remains relatively un-affected in the mutants but the processing of olfactory information in the central brain is rendered defective, resulting in an olfactory CO_2_-induced irreversible wing hold up phenotype. Overexpression of either Drosophila Prl-1 or hPRL in the nervous system could rescue the mutant phenotype. The ablation of the olfactory CO_2_ sensory neurons blocks phenotype development. Therefor CO_2_ might act as a neuropathological substrate for brain disorders in absence of Prl-1.

CO_2_ has also had a long history as a human anesthetic where increased CO_2_ levels decreases hippocampal neuronal excitability and may cause sedation (Capps, 1968, Jones, 1984). At the cellular level, CO_2_-induces changes in pH and may alter neuronal excitability by altering extracellular adenosine and ATP concentrations (Xu, Uh et al., 2011). CO_2_ insufflation-induces oxidative stress and has a toxic effect on neuroblastoma cells leading to DNA damage (Montalto, Currò et al., 2013). In *E. coli*, CO_2_ was seen to exacerbate the toxicity of ROS in a dose-dependent manner (Ezraty, Ducret et al., 2011). Here, our observation that *prl-1* mutants display both enhancement of Ca^2+^ transients in the CO_2_-responsive neural circuit and elevated ROS production and that both could be rescued by overexpression of Prl-1 in the nervous system, suggests a potent neuropretective role for Prl-1. We propose that CO_2_ could act as an olfactory pathway linked neuropathological substrate for brain disorders in absence of Prl-1. This finding might open up a new focus upon the functional pathway of the hPrl beyond studies of its oncogenic properties.

It is notable that, in *prl-1* mutants, the expression of Uex is clearly decreased. Knockdown of Uex resulted in the same wing hold up as observed in the *prl-1* mutants, while abnormal wing posture in *prl-1* mutants could be rescued by the expression of Uex in the nervous system. Thus irreversible wing hold up could be regarded as a downstream consequence of a deficiency in Uex due to the absence of Prl-1. The mammalian counterparts of Uex have been predicted as magnesium transporters involved in tumor progression (Gulerez, Funato et al., 2016, Kostantin, Hardy et al., 2016). Biochemical analyses of cultured cells revealed that the PRL acts through the inhibition of CNNM to regulate intracellular magnesium homeostasis (Funato et al., 2014, Hardy et al., 2015, Hirata, Funato et al., 2014). In vertebrate neurons, Mg^2+^ acts as a physiological Ca^2+^ antagonist for blocking the excitatory N-methyl-D-aspartate receptors in the CNS (Iseri & French, 1984, Zito & Scheuss, 2009) and has therefore been suggested as a possible means of resolving muscle rigidity and spasm in cases of tetanus(Ceneviva, Thomas et al., 2003). In our experiment, although it was inaccessible to measure the Mg^2+^ status in the prl-1 mutants, enhanced Ca^2+^ activities were observed at CO_2_-responsive neuron synaptic terminals. In human, CNNM2 mutations cause impaired brain development and seizures in patients with hypomagnesemia (Arjona et al., 2014). We hypothesize that the decreased Uex in Prl-1 mutants could cause disturbed Mg^2+^ homeostasis and consequently lead to a brain disorder presented as irreversible wing hold up.

The irreversible wing hold up in *prl-1* mutants seems to be a prominent form of hyperreflexia resembling spasticity. Spasticity in mammals is a common complication beyond brain injury, presenting as intermittent or sustained involuntary activation of muscles (Pandyan, Gregoric et al., 2005). In an early study Takano *et al* had noted that mPRL is expressed in neurons and oligodendrocytes in the brain and is enhanced in the cerebral cortex following transient forebrain ischemia in rats (Takano, Fukuyama et al.,1996). The possible role of PRL-1 in an ischemia stroke situation where neurons undergo restrictions in both oxygenated blood inflow and CO_2_ blood-export, has not yet to be followed up. Analysis of the cerebral cortex and hippocampus tissue sections from a small sample set of Alzheimer’s disease patients revealed that there seemed a trend toward increased expression of PRL-1 mRNA expression and also a considerable correlations with patient age in the brain (Dumaual et al., 2012). The *prl-1* mutant flies also exhibited age-dependent phenotypic manifestation. Intriguingly, in Parkinson’s and Alzheimer’s diseases there is profound olfactory disorder in odor threshold detection, odor memory, and/or odor identification occurring prior to disease onset (Doty, 2017, Kalia & Lang, 2015, Rudzinski, Fletcher et al., 2008), often associated with aspects of limb spasticity (Erro & Stamelou, 2017, Karlstrom, Brooks William et al., 2007). *Drosophila* models of human neurodegenerative diseases have already included observations of the wing hold up or hold out phenotypes (Clark, Dodson et al., 2006, Freibaum, Lu et al., 2015, Pandey & Nichols, 2011).The cause of these disorders is unclear. It is worth considering that olfactory CO_2_ stimulation, even of atmospheric levels, might be directly involved in these neurologic disease-related detrimental processes, for which Prl-1 provides defense. Our findings might be applicable for the potential involvement of hPRL in neuroprotection.

## Materials and methods

### Fly strains and genetics

The following transgenic flies were used: (1) *Elav-Gal4*, (2) *Tubulin-Gal4*, (3) *Repo-Gal4*, (4) *TH-Gal4*, (5) *Orco-Gal4*, (6) *Or476-Gal4*, (7) *Gr21a-Gal4*, (8) *Gr63a-Gal4*, (9) *V201089-Gal4*, (10) *V200516-Gal4*, (11) *MB247-Gal4*, (12) *OK107-Gal4*, (13) *GF-Gal4*, (14) *D42-Gal4*, (15) *24B-Gal4*, (16) *Vglut-Gal4*, (17) *MHC-Gal4*, (18) *Mef2-Gal4*, (19) *Elav-GeneSwitch-Gal4*, (20) *Uas-mCD8::GFP*, (21) *Uas-GCaMP6.0*,(22) *Uas-uex IR*, (23) *Uas-Gr21a IR*, (24) *GSTD-ARE-GFP*.

Plasmid construction and infection: *prl-1* was cloned in a *pAHW* vector (Carnegie Institution of Washington), and *pGEX-4T-1* vector. Mutations were introduced by site-directed mutagenesis using a GBclonart mutagenesis kit.

Generation of transgenic flies: *prl-1*, *EGFP-prl-1*, *uex*, h*Prl-1*, and h*Prl-2* were cloned in a *pUAST-attb* plasmid using specific primers (see below) and amplified by PCR. The *prl-Gal4* was cloned in a *pW25-Gal4* plasmid, which flanked the *Gal4* sequence with both the 5’ and 3’ flanking regions of the *prl-1* gene. All constructs were integrated into a single *attP* docking site, *VK33* on chromosome 3L, using common *phiC31* site-specific integration, as Matthew P Fish described(Venken & Bellen, 2007).

Antibody generation: Full-length cDNA of *prl-1*, and N-terminal 300bp cDNA of *uex* were cloned into a pGEX-4T-1 expression vector. The GST-fusion protein was affinity purified using Sepharose-4B beads (GE Healthcare). The polyclonal antibody was raised in rabbits. Animal experiments were conducted in accordance with the Guidelines for the Care and Use of Laboratory Animals of Zhejiang University.

### Immunofluorescence staining

Immunofluorescence staining of the adult brains and S2 cells were conducted as previously described (Riemensperger, Strauss et al., 2011, Rogers, Wiedemann et al., 2004). The following primary antibodies were used: rabbit anti-Prl-1,1:500 (this study); rabbit anti-Uex,1:500; mouse anti-Brp^nc82^, 1:50 (DSHB);DAPI (1 g/ml; Sigma-Aldrich).

### IP and GST pull-down

Cells and fly tissues were lysed with TAP buffer (1% Triton, 50mM Tris pH 8.0, 125mM NaCl, 5% Glycerol, 0.4% NP-40, 1.5mM MgCl_2_, 1mM EDTA, 25mM NaF, and 1mM Na_3_VO_4_), supplemented with protease inhibitor (Roche, Laval, QC, Canada). For IP, 1-2 mg of proteins was incubated with 1 μg of HA antibody (Abcam, ab9110) and Protein A-agarose beads (Roche Applied Science) according to the manufacturer’s protocol. The supernatants eluted from immunoprecipitated beads were loaded for Western blotting following standard protocols. For the GST pull-down assays, 500 μg of proteins were incubated with glutathione sepharose (GE Healthcare, Canada) for 3 h.

### Calcium imaging

Sample preparation and calcium imaging were as described in. CO_2_ was delivered at a flow rate of 5 ml min^−1^. Sample preparation and calcium imaging were as described in(Jones et al., 2007, Suh et al., 2004). CO_2_ was delivered at a flow rate of 5 ml min^−1^. The adult *Drosophila* was fixed to the scotch tape by its dorsal parts with its wings and the maxillary pulp was also immobilized using a scotch tape slice. The dorsal parts of the adult brain were dissected within a drop of AHL solution(Jones et al., 2007, Suh et al., 2004) and then covered with a coverslip for Ca^2+^ imaging. Imaging of Ca^2+^ was performed on Olympus fluoresce microscope with a x20 objective lens. Images were acquired at 1.42 frames per second. For quantitative analysis of Ca^2+^ imaging data, images were batch processed with Image J to determine fluorescence intensity. The initial 120 seconds sequential images, occurring prior to the 20 second CO_2_ stimulus, were subjectively selected and the average fluorescence intensity (F) was set as the basal level. Changes in fluorescence intensity (△F) in the images were calculated and △F/F was used to denote Ca^2+^ responses. Heat map images were generated with Matlab (Mathworks Inc., Natick, MA, USA) by setting the basal fluorescence level at zero.

### Behavior assays

About 50 flies were placed into the T-maze for each CO_2_ avoidance assay. At the T-maze choice point, flies sensed the converging currents of fresh air and CO_2_ from each arm. After 1 min of choice, the avoidance index (AI) was calculated.

### Statistics

All the raw data were analyzed parametrically with excel and GraphpadPrism 5 software. The data were evaluated via Two-tailed Student’s t test. All data are presented as mean ±SEM. \**P* < 0.05; \*\**P* < 0.01; \*\*\**P* < 0.001.

## Acknowledgments

We thank Jun Ma, Hao Wang, Lijun Kang for helpful suggestions on data analysis; Qi Zeng, Chris Wood and Jingwei Zhao for discussions and comments on the manuscript. We also thank Yu Cai (GSTD1 - ARE-GFP), Jiangqu Liu (GCaMP), the Bloomington Drosophila Stock Center and Qinghua Drosophila Stock Center for providing the fly stocks. This work was supported by the National Basic Research Program of China (2013CB945600 (X.Y.), 2012CB966800 (W.G.)).

---

**Fig. S1.**
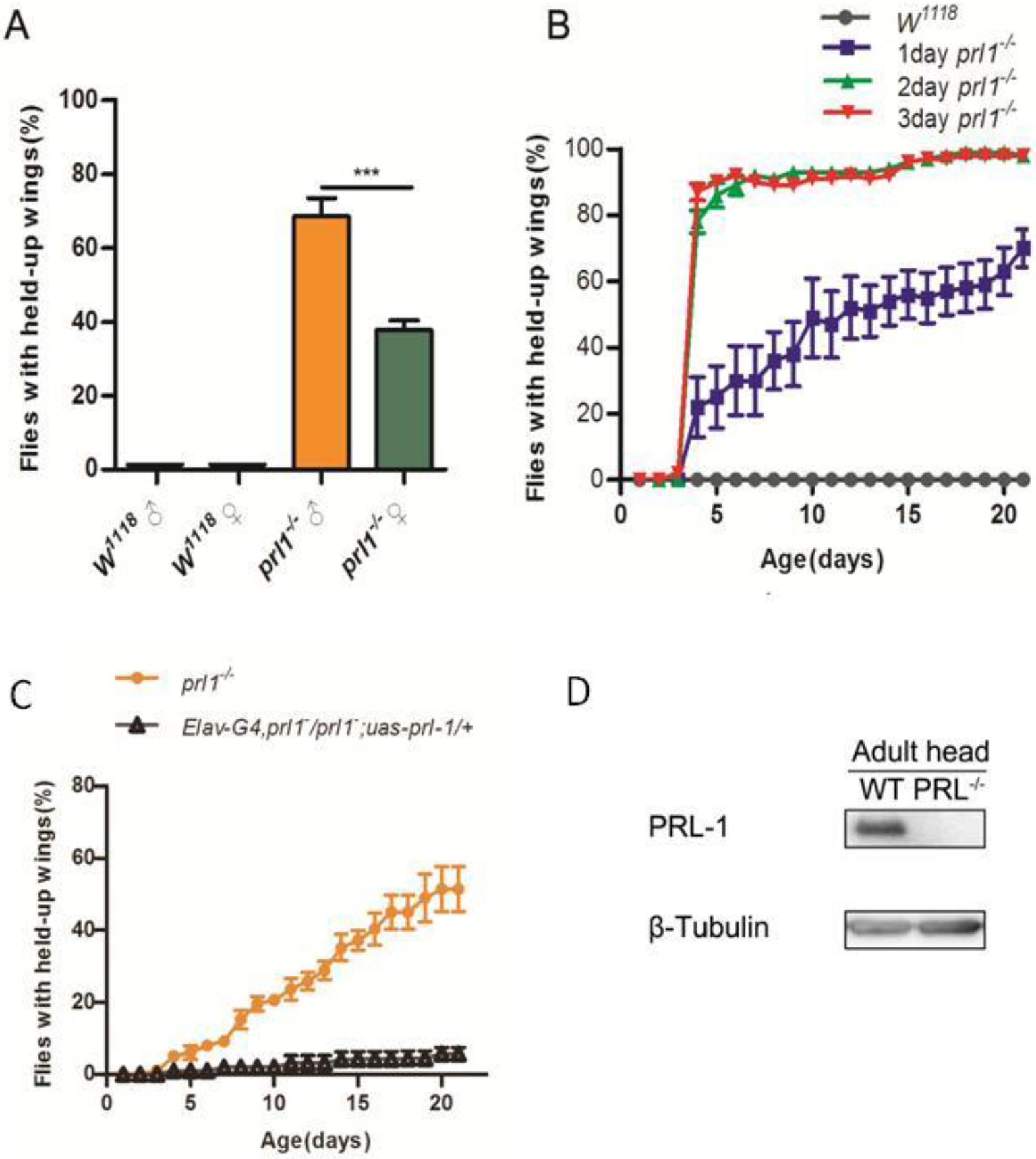
Ocurrences of wing hold up phenotype in absence of Prl-1 and expression of Prl-1 in adult brains. **(A)** Male and female 3-day old mutants and wild type flies (n=100 for each group) observed 12 hours after recovery from acute CO_2_ exposure. *prl-1* male mutant animals displayed a more prevalent wing hold up than the female (over 60% as compared to under 40%). **(B)** The 1-day, 2-day, and 3-day old male mutants treated with acute CO_2_ and flies with irreversible holding up of wingswere counted for 21 days. Results are plotted as means ± SEM. Two-tailed Student’s t test with ***p*0.001, n=180 for each group. **(C)** Elav-Gal4 could rescue the wing hold up of *prl-1* mutants in the ambient environment. Genotype: *Elav-Gal4, prl-1^−^/prl-1^−^; uas-prl-1/+* (n=100 per group). **(D)** Western blot on adult brains extracted from wild type and *prl-1* mutant animals. Lysates were probed with anti-Prl-1 and anti-Tubulin antibodies.

**Fig. S2.**
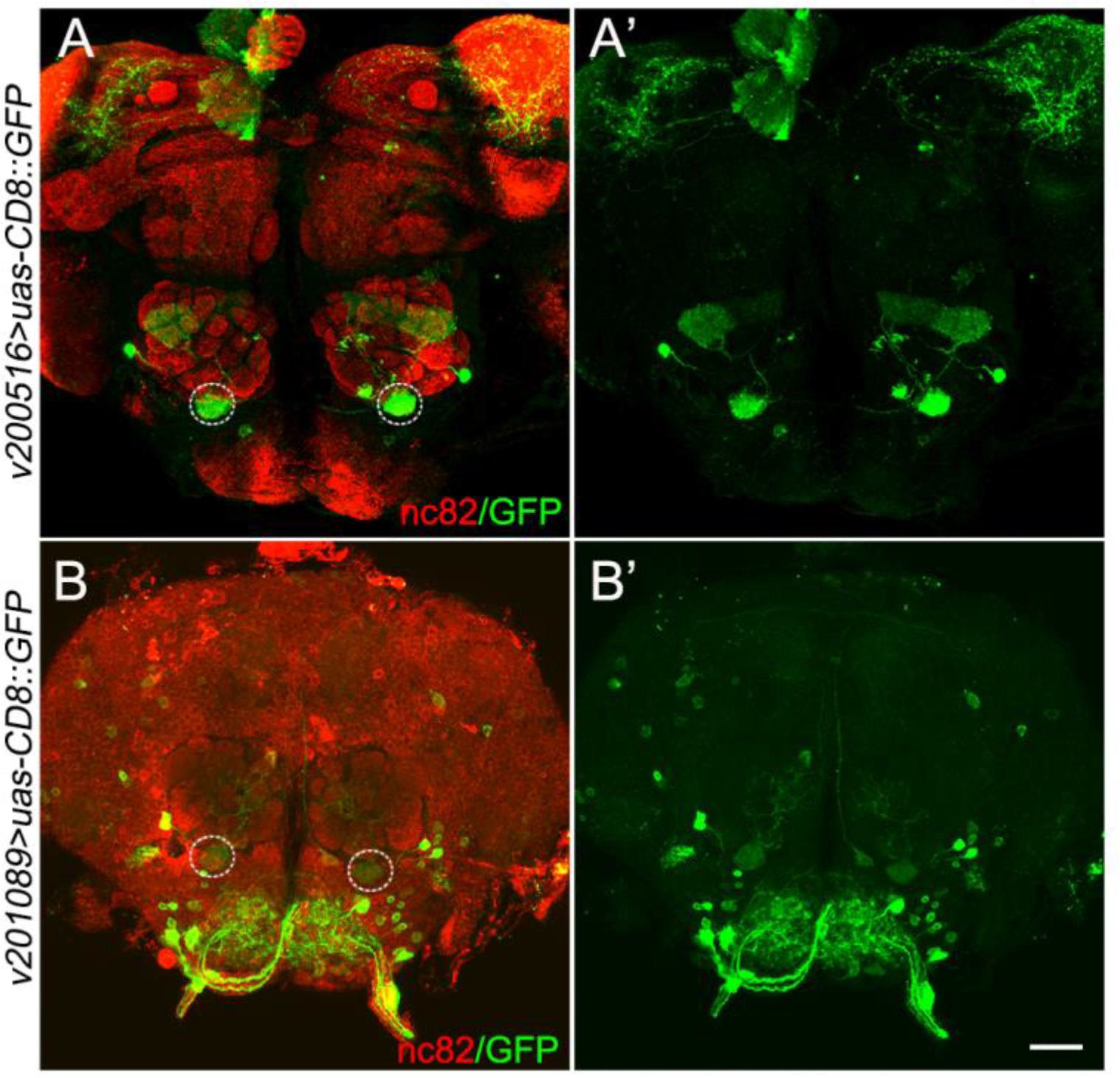
CO_2_ responsive projection neurons in the stereotyped V-glomeruli in the antennal lobe. Expression of two *PNv-Gal4*s (v201089, v200516) in V-glomeruli (dotted circle). Gal4 neurons were labeled with *UAS-mCD8::GFP*(green), and brain neuropils were immunostained with a Brp^nc82^ antibody (red). Scale bar: 30um.

**Fig. S3.**
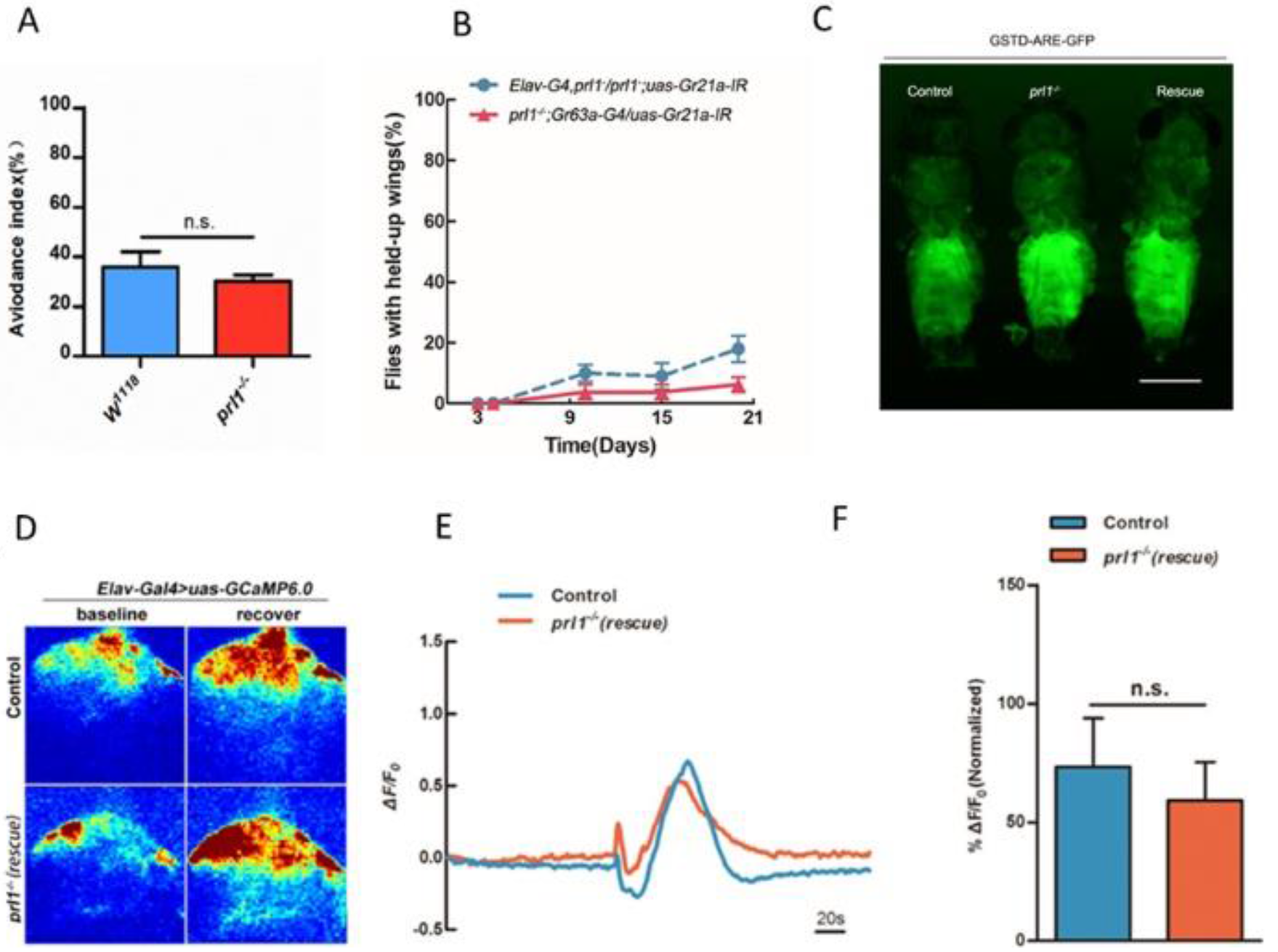
Enhancement of CO_2_ sensitivity and ROS production in *prl-1* mutants can be rescued by ectopic expression of Prl-1 in the nervous system. (A) Analysis of CO_2_-evoked avoidance responses. Compared with wild type flies, *prl-1* mutants exhibit no significant difference in CO_2_ avoidance behavior in a T-maze test.Results are plotted as means ± SEM. Two-tailed Student’s t test with p>0.05, n=40-50 for each group. (B) Ablation of Gr21a receptor suppresses the wing hold up. In *prl-1* mutant background, genetic ablation of CO_2_ sensory neurons driven by *Elav-Gal4* or *Gr63a-Gal4* could inhibit the development of wing hold up for flies in the ambient environment. n=100 for each group. (C): Activation of the *GSTD1* enhancer in *prl-1* mutants. The activity of *GSTD1* in the *prl-1* mutants was reduced by overexpression of Uas-Prl-1 with Elav-Gal4 driver. Genotype: control: *GSTD-GFP/+*;*prl-1* mutants: *GSTD-GFP, prl1^−^/prl1^−^* Rescue: *GSTD-GFP, prl1^−^/Elav-Gal4,prl1^−^;uas-prl-1/+*. Scale bar: 200x pix. (n=120 each group). (D-F): Calcium responses in the antennal lobe with CO_2_ stimulation for control or *prl-1* rescued flies (D). Representative images of GCaMP6.0 fluorescence (%ΔF/F) obtained from in vivo imaging at baseline and after CO_2_ stimuli to control or *prl-1* rescue flies (E). GCaMP6.0 responses from control or *prl-1* rescue flies across CO_2_ presentation with bar graph quantitation. Statistics for (E) results are plotted as means ± SEM (F). Two-tailed Student’s t test with p>0.05, n=4-7 for each group.

**Fig. S4.**
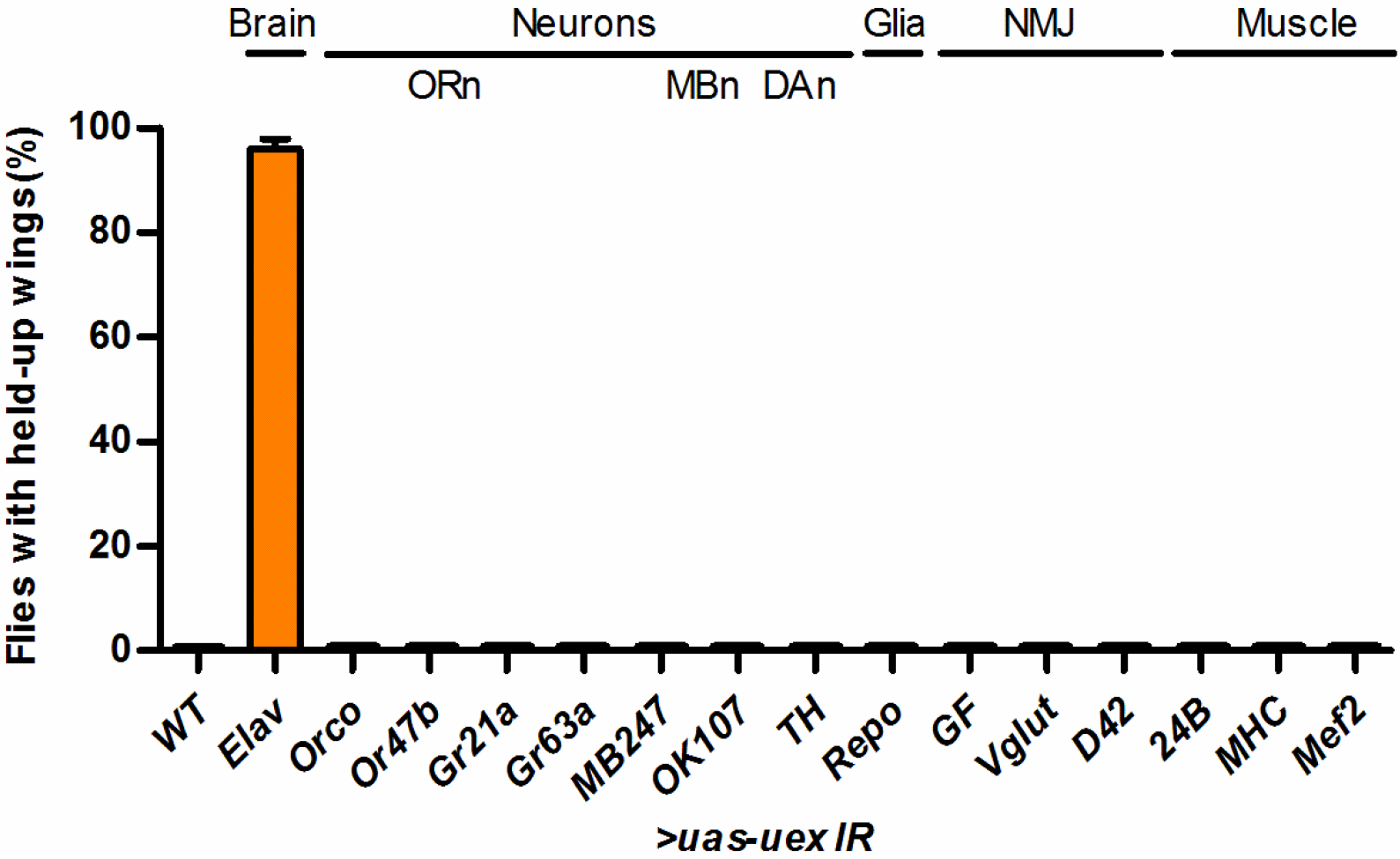
Wing hold up phenotype induced by knockdown of Uex in the brain. RNAi target of the *uex* gene was tested along with a battery of *Gal4* lines: *Elav-Gal4* (pan-neuronal), *Orco-Gal4*, *Or47b-Gal4*, *Gr21a-Gal4* and *Gr63a-Gal4* (olfactory receptor neurons), *MB247-Gal4* and *OK107-Gal4* (MBneuron), *TH-Gal4* (DA neuron),*Repo-Gal4* (glia), *GF-Gal4* (giant fiber), *Vglut-Gal4* (Vglut), *D42-Gal4* (NMJ), *24B-Gal4* (muscle), *MHC-Gal4* (muscle), and *Mef2-Gal4* (muscle).(n=60 each group)

Primer lists:

*prl-1-F*: caagaagagaactctgaataATGAGCATCACCATGCGTC
*prl-1-R*: aggttccttcacaaagatccCTATTGCACAGAACATGAATTC
*hPrl-1-F*: tacgctgctcatggcggaATGGCTCGAATGAACCGCC
*hPrl-1-R*: aggttccttcacaaagatccTTATTGAATGCAACAGTTGT
*hPrl-2-F*: tacgctgctcatggcggaATGAACCGTCCAGCCCCT
*hPrl-2-R*: aggttccttcacaaagatccCTACTGAACACAGCAATGCC
*EGFP-prl-1-F*: attcgttaacagatctgcATGGTGAGCAAGGGCGAGG
*EGFP-prl-1-R*: tcacaaagatcctctagagCTATTGCACAGAACATGAAT
*EGFP-prl −1-F1*: gcatggacgagctgtacaAGATGAGCATCACCATGCGTC
*EGFP-prl −1-R1*: gacgcatggtgatgctcaTCTTGTACAGCTCGTCCATGC
*uex-F*: agaagagaactctgaataATGAACACATATTTCATATC
*uex-R*: ttccttcacaaagatccTTAGGGCTTACTTTGCTTGCTCT
*prl-Gal4-F*: aattgggaattcgttaacaTCACCATCCGTGTCTACCAAC
*prl-Gal4-R*: atctttcaggaggcgcggccACAATTACAAAAGCTGTTCT
*HA-prl-1-F*: tacgctgctcatggcggaATGAGCATCACCATGCGTC
*HA-prl-1-R*: agaagtaaggttccttcaca CTATTGCACAGAACATGAATTC
*BamHI-prl-1-F*: CGggatccATGAGCATCACCATGCGTC
*NotI-prl-1-R*: TTgcggccgcTTGCACAGAACATGAATTC
*BamHI-uex-F*: CGggatccGCAAAAAAAGCTACATTG
*NotI-uex-R*: TTgcggccgcCATGGAAGCAGCTGTCGT
*D77A- prl-1-F*: GCCTTTGAGGcCGGCACCT
*D77A- prl-1-R*: AGGTGCCGgCCTCAAAGGC
*C109S- prl-1-F*: GCCGTTCATaGTGTGGCTG
*C109S- prl-1-R*: CAGCCACACtATGAACGGC
*prl-1-gRNA target*: GGTTATGTCTGATGGTCGATcgg
*uex-gRNA target*: GGCACTCCGAGTCGCCCAGagg

